# Natural variation in *Arabidopsis thaliana* highlights a key role of Glyoxalase I;2 in detoxifying glucose-derived reactive carbonyl species

**DOI:** 10.1101/2023.12.20.572487

**Authors:** Maroua Bouzid, Manuel Balparda, Aylin Kerim, Ying Fu, Clarisa E. Alvarez, Saleh Alseekh, Alisdair R. Fernie, Veronica G. Maurino

## Abstract

Reactive carbonyl species (RCS) are toxic byproducts of normal metabolism that become more prevalent under oxidative stress. Here, we show that *Arabidopsis thaliana* ecotypes exhibit natural variation in their ability to detoxify glucose-derived RCS. We identified the IP-Pal-0 ecotype showing enhanced tolerance to glucose-derived RCS via upregulation of the glyoxalase system. In particular, the Viridiplantae-specific isoforms GLXI;2, GLXII;4, and GLXII;5 are highly expressed in IP-Pal-0 when plants are grown in the presence of 2-keto-D-glucose (KDG; glucosone) or methylglyoxal, and protein extracts from these plants display enhanced GLXI activity on KDG than other ecotypes. We identified specific motif/*cis*-regulatory elements in the *GLXI;2* promoter regions of Col-0 and IP-Pal-0 that may underlie the differential expression of *GLXI;2* associated with KDG detoxification. IP-Pal-0 GLXI;2 contains two different amino acids compared to Col-0, but these do not affect the basic kinetics of the protein. Interestingly, we found that the simultaneous change of these amino acids also occurs together in the GLXI proteins of some other organisms, suggesting a convergence in the simultaneous change of both amino acid residues. Our findings underscore the importance of Viridiplantae-specific glyoxalase isoforms in detoxifying glucose-derived RCS, particularly KDG, and highlight the promise of harnessing natural genetic diversity in the glyoxalase pathway to enhance plant stress tolerance.

## Introduction

The study of natural plant ecotypes continues to unveil critical insights into the biological mechanisms underlying environmental adaptation. By studying genes associated with stress responses, it is possible to identify those that are key to resilience in natural variants of *Arabidopsis thaliana*. Various ecotypes of this model plant have been linked to tolerance against cold (1), drought (2), heat (1), and even viral infections (3).

Environmental stresses and normal physiological processes can both lead to the accumulation of reactive small molecules, which can compromise plant fitness (4). Two well-known examples are reactive oxygen species (ROS) and reactive carbonyl species (RCS), both of which threaten cellular integrity by reacting with macromolecules (5, 6). Common RCS in biological systems include methylglyoxal (MGO), glyoxal (GO), 2-keto-D-glucose (KDG; glucosone), and 3-deoxyglucosone (7, 8). These compounds arise mainly as byproducts of sugar metabolism through glucose oxidation (9) and glycated protein cleavage (10). Various abiotic stresses have been associated with increased RCS accumulation, including salinity stress in *Cucurbita maxima* (11), drought stress in *Vigna radiata* and *Brassica juncea* (12, 13), and exposure to toxic metals (14).

To mitigate the toxic effects of RCS, plants have evolved enzymatic detoxification mechanisms, the most prominent being the glyoxalase (GLX) system (4). This system consists of two enzymes: glyoxalase I (GLXI), an S-D-lactoylglutathione lyase, and glyoxalase II (GLXII), an S-2-hydroxyacylglutathione hydrolase. Together, they catalyze the conversion of harmful RCS into less toxic metabolites using glutathione (GSH) as a cofactor (8, 15, 16).

GLXI is a lyase that depends on a divalent metal ion as cofactor. It uses as substrate a hemithioacetal formed by the non-enzymatic reaction of an RCS with glutathione (GSH), resulting in the formation of a glutathione thioester in which GSH is covalently bound to the RCS (17). GLXII then catalyses the conversion of the glutathione thioester to a (D)-α-hydroxy acid derivative which can be reintegrated into cellular metabolism, releasing GSH. This pathway enables the conversion of MGO into D-lactate, which can be further metabolized into pyruvate by D-lactate dehydrogenase (18, 19). Additionally, GLXI;2 has been shown to act on KDG, into D-gluconic acid (8).

In *A. thaliana*, three gene loci encode functional GLXI isoforms: *GLXI;*1 (At1g67280), *GLXI;2* (At1g11840), and *GLXI;3* (At1g01180) (15). These isoforms belong to two evolutionarily distinct groups. The first group, including GLXI;1 and GLXI;2, is specific to Viridiplantae and bacteria (8). These monomeric enzymes contain two structural domains, one forming an active site and the other a cryptic site of unknown function (8, 20–22). They are classified as Ni²⁺-type GLXI, preferring Ni²⁺ when MGO-GSH is the substrate and Mn²⁺ when GO-GSH is the substrate (15). The second group, represented by GLXI;3, is conserved across eukaryotic kingdoms (8, 23). These isoforms form homodimers with two functionally equivalent active sites in Animalia and Viridiplantae, whereas other groups such as Rhodophyta, Phaeophyta, Diatoms, Protozoa, and Fungi exhibit both single- and two-domain forms (8). *A. thaliana* GLXI;3 is classified as a Zn^2+^-type GLXI but has a strong catalytic preference for Mn²⁺ (15, 20, 22).

Similarly, three gene loci encode GLXII homologs: GLXII;2 (At3g10850), GLXII;4 (At1g06130), and GLXII;5 (At2g31350) (15). These enzymes possess a Fe^3+^/Zn^2+^ binuclear metal center and operate as dimers, catalyzing reactions with a wide range of glutathione thioesters (16, 22, 24, 25). As with GLXI, plant GLXII isoforms have distinct evolutionary origins: while GLXII;2 is broadly conserved across eukaryotic kingdoms, GLXII;4 and GLXII;5 are exclusive to Viridiplantae and Proteobacteria (8).

The glyoxalase system has been extensively studied for its role in enhancing stress tolerance in crops. Overexpression of GLX components in transgenic plants has led to improved resistance to abiotic and biotic stresses, including enhanced salt tolerance in tobacco and brown mustard (13, 26), and increased resistance to drought, heat, salinity, and pathogens in transgenic rice (27–29). However, regulatory restrictions, such as EU Directive 2015/412, which limits the commercial use of genetically modified crops in Europe, highlight the importance of exploring alternative approaches (https://eur-lex.europa.eu/eli/dir/2015/412/oj). Harnessing natural genetic variation within the GLX system across *A. thaliana* ecotypes may provide a promising, non-GMO strategy for improving crop resilience (30, 31).

*A. thaliana* ecotypes thrive in diverse environments and exhibit remarkable phenotypic plasticity in response to variations in light, temperature, humidity, water availability, and soil composition (32). Numerous studies have highlighted significant genetic and physiological differences among ecotypes (33–35). Investigating the functional diversity of the GLX system across these ecotypes could help identify genetic determinants of stress resilience.

In this study, we analyzed *A. thaliana* ecotypes to identify those with enhanced RCS detoxification capacity by assessing their growth performance under MGO and KDG exposure. Among them, IP-Pal-0 showed significantly enhanced growth compared to the reference ecotype Col-0 in both MGO- and KDG-containing media. Molecular and physiological analyses revealed that the expression of the Viridiplantae-specific GLXI;2 isoform is associated with the observed differences in stress tolerance between ecotypes. These findings underscore the potential of harnessing natural genetic variation in the glyoxalase system to improve plant resilience to stress.

## Methods

### Growth conditions

*A. thaliana* ecotypes listed in Figure 1 were kindly provided by Prof. Juliette de Maux (University of Cologne). All seedlings were grown either on soil or sterile media under long-day conditions (16 hours light/8 hours dark) at 22 °C and 55% relative humidity. Illumination was provided by a combination of Osram HO 80W/840 and HO 80W/830 fluorescence bulbs, delivering a light intensity of 120 μE^-2^ s^-1^.

**Figure 1.**
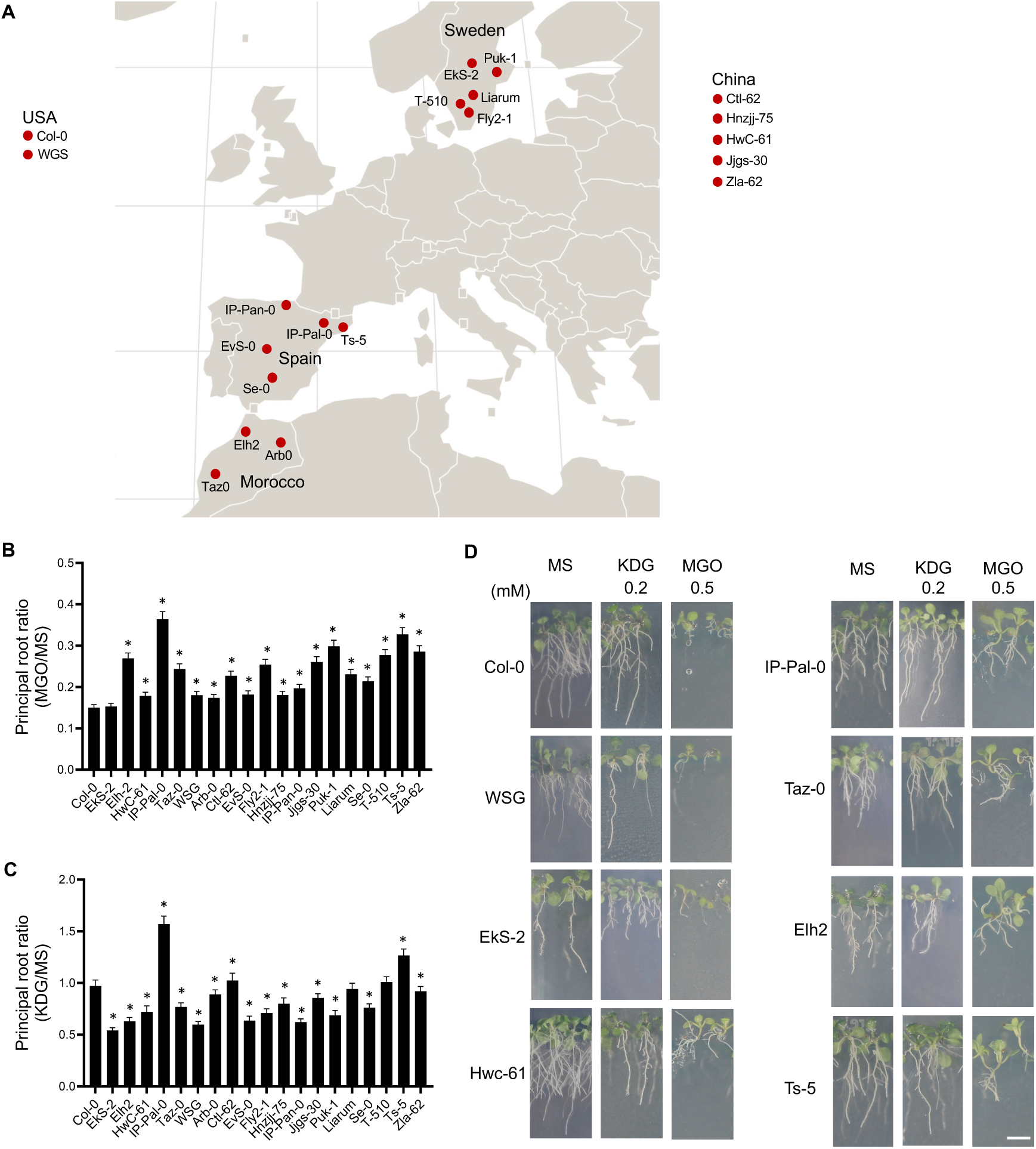
Effect of KDG and MGO on the development of *A. thaliana* ecotypes. (A) Geographical origin of the ecotypes analyzed. For those from Spain and Sweden, the localisation shows the exact coordinates provided by 1001genomes (https://1001genomes.org). (B) Main root length after 14 days of germination on media containing 0.5 mM MGO relative to the length in control media (MS). (C) Main root length after 14 days of germination on media containing 0.2 mM KDG relative to the length in MS. The graphs show the means and standard errors of the measurements of 3 independent experiments (*n*=3) of 20 roots each. Significant differences were analyzed by one-way ANOVA. An asterisk (*) indicates a significant difference (P<0.001) between an ecotype and Col-0 by Dunnett’s multiple comparison test. (D) Images of seedling of selected ecotypes at 14 d after germination in different growth media. Scale bar = 1 cm.

Sterilized seeds of each *A. thaliana* accession were sown on plates with 0.5X MS (Duchefa), either alone or supplemented with 0.2 mM KDG (Sigma-Aldrich, USA) or 0.5 mM MGO (Sigma Aldrich, USA), and vernalized at 4 °C in darkness for 48 hours. Additional growth experiments were performed on 0.5X MS medium supplemented with either 2% or 4% glucose. For each ecotype and treatment, 15 seeds were aligned in a single row per plate. Following vernalization, plates were positioned vertically in a growth cabinet (Percival Scientific Inc., USA) under long-day conditions. Primary root length was measured for each seedling using the ImageJ software (36).

### RNA extraction and cDNA synthesis

Total RNA was isolated from roots and shoots of plants cultivated for 14 days in 0.5X MS medium, either alone or supplemented with 0.2 mM KDG. RNA from rosette leaves was extracted using the Universal RNA Purification Kit (Roboklon GmbH, Germany), following the manufacturer’s protocol. Elution was performed in 30 µl of RNase-free distilled water. RNA quantity and purity were assessed using a NanoDrop Nabi™ UV/Vis spectrophotometer (MicroDigital Co. Ltd, South Korea) and confirmed by agarose gel electrophoresis. The extracted RNA was stored at −20 °C until further processing.

To remove residual genomic DNA, RNA samples were treated with the DNA-free kit (Ambion, Thermo Fisher Scientific, USA). Subsequently, 2 µg of RNA was reverse transcribed into complementary DNA (cDNA) using the RevertAid First Strand cDNA Synthesis Kit (Thermo Fisher Scientific, USA) with oligo(dT) primers, according to the manufacturer’s instructions. Briefly, RNA was mixed with 1 µl of oligo(dT) primer and incubated at 65 °C for 10 minutes. The reaction mix was then supplemented with 2 µl dNTPs, 0.8 µl RevertAid H Minus M-MuLV Reverse Transcriptase, 4 µl of 5X reaction buffer, and 10 µl of RNase-free water. Reverse transcription was carried out at 42 °C for 60 minutes, followed by enzyme inactivation at 70 °C for 10 minutes. The resulting cDNA was stored at −20 °C for subsequent use.

### Reverse-transcription quantitative PCR

Reverse-transcription quantitative PCR (qRT-PCR) was performed using the KAPA SYBR FAST qPCR kit (KAPA Biosystems Inc., Switzerland) to assess the expression levels of *GLXI;1*, *GLXI;2*, *GLXI;3, GLXII;2, GLXII;4 and GLXII;5* in the cDNA samples from various *A. thaliana* ecotypes. Reactions were run on a Mastercycler® ep realplex system (Eppendorf, Germany), and cycle threshold (Ct) values were obtained using the associated realplex software. Expression levels were normalized to the housekeeping gene *Actin2* (ACT2), which served as the internal control.

### Protein Isolation from plant tissue and GLXI enzymatic assay

Fourteen-day-old seedlings cultivated on 0.5X MS medium, with or without 0.2 mM KDG, were harvested and stored at −80 °C until analysis. For protein extraction, 200 mg of frozen plant tissue was ground in liquid nitrogen and suspended in extraction buffer containing 100 mM Tris-HCl (pH 8.0), 100 mM NaCl, 0.5% (v/v) Triton X-100, 2 mM PMSF, and 1% (w/v) PVP40. The mixture was centrifuged at 20,000 × g for 10 minutes at 4 °C, and the resulting supernatant was used for enzymatic activity assays.

To evaluate total GLXI activity, 2-keto-D-glucose (KDG; Sigma-Aldrich, USA) was pre-incubated with 1 mM reduced glutathione (GSH) for 10 minutes to form the hemithioacetal substrate. Reaction mixtures were set up with increasing concentrations of KDG (0, 0.25, 0.5, 1, and 2 mM), while the GSH concentration was kept constant at 1 mM. Each reaction received 20 μg of protein extract and was incubated at 25 °C for 30 minutes. Absorbance at 412 nm was recorded, after which 1 mM 5,5′-dithiobis-(2-nitrobenzoic acid) (DTNB; Geno Technologies Inc., USA) was added to quantify unreacted GSH. A second absorbance measurement was taken at 412 nm following a 10-minute incubation. All components were prepared in 0.1 M MOPS buffer (pH 7.0). Measurements were performed using a Spark Multimode Microplate Reader (Tecan, Germany). A standard curve generated from 0.2-2 mM GSH was used to determine GSH concentrations. Each assay included a minimum of three biological and three technical replicates.

### Heterologous expression and purification of GLXI;2 variants

The pET16b expression constructs encoding GLXI;2 variants (*GLXI;2*-Phe278 and *GLXI;2*- Ile278) were transformed into *E. coli* Rosetta2 (DE3) pLysS cells. Transformed cultures were grown in 400 ml LB medium at 37 °C with continuous shaking until reaching an OD₆₀₀ of 0.4-0.6. ampicillin (50 μg/ml) and chloramphenicol (34 μg/ml) were used for selection. Protein expression was induced with 1 mM isopropyl β-D-1-thiogalactopyranoside (IPTG), followed by a 4-hour incubation at 37 °C with shaking. Cells were harvested by centrifugation at 6,000 × g for 15 min and pellets were stored at −20 °C until further processing.

Frozen pellets were thawed on ice and resuspended in 20 mM Tris-HCl (pH 8.0) containing 500 mM NaCl, 5 mM imidazole, 1 mM phenylmethylsulfonyl fluoride (PMSF), and a spatula-tip of lysozyme. The suspension was sonicated and centrifuged at 14,000 × g for 20 min at 4 °C. The resulting supernatant was subjected to immobilized metal ion affinity chromatography using Ni-NTA agarose (Thermo Fisher Scientific, USA) under gravity flow.

The Ni-NTA column was pre-equilibrated with 20 mM Tris-HCl (pH 8.0) containing 500 mM NaCl and 5 mM imidazole. After loading the supernatant, the column was sequentially washed four times with the same buffer containing increasing concentrations of imidazole (5, 40, and 60 mM). Proteins were eluted with 1 ml of buffer containing 20 mM Tris-HCl (pH 8.0), 500 mM NaCl, and 200 mM imidazole. Eluted proteins were concentrated and buffer-exchanged into 20 mM Tris-HCl (pH 7.5) using Vivaspin 30K centrifugal concentrators (Sartorius, Göttingen, Germany).

### Native polyacrylamide gel electrophoresis

Native-PAGE was performed using 10% (w/v) polyacrylamide gels, following the method described by Laemmli (37). Protein separation was carried out alongside the SERVA Marker Mix for Blue/Clear Native PAGE (SERVA Electrophoresis GmbH, Heidelberg, Germany). Gels were stained with Coomassie Brilliant Blue to visualize protein bands.

### Determination of kinetic constants

GLXI enzymatic activity was measured spectrophotometrically using a Spark® microplate reader (Tecan) in 96-well UV-Star Microplates (Greiner Bio-One). The standard assay buffer consisted of 100 mM MOPS (pH 7.0), 3.5 mM methylglyoxal (MGO) or glyoxal (GO), 1.7 mM reduced glutathione (GSH), and 0.5 mM of the appropriate divalent metal ion, Ni²⁺ for MGO and Mn²⁺ for GO (15). The assay relies on the spontaneous formation of hemithioacetal (HTA) through the non-enzymatic reaction between the reactive carbonyl species and GSH, which serves as the natural substrate for GLXI enzymes. Kinetic parameters were determined following the protocol described by Schmitz et al. (15). All measurements were conducted in at least three independent biological replicates.

### Homology modeling of GLXI;2 3D structure

The three-dimensional structure of *A. thaliana* GLXI;2 was predicted using the AlphaFold-based ColabFold server (38) with the primary amino acid sequence as input. Structural visualization and measurement of distances between amino acid residues were performed using PyMOL version 2.5.4 (Schrödinger; https://pymol.org/).

### Metabolite abundance analysis by gas chromatography-mass spectrometry

Whole seedlings were harvested after 14 days of growth on 0.5X MS medium or 0.2 mM KDG-supplemented medium for gas chromatography-mass spectrometry (GC-MS) analysis. Metabolite extraction, derivatization, and GC-MS analysis were performed following the protocol described by Lisec et al. (39), using the same equipment configuration and settings. Shortly, 200 μl of chloroform and 300 μl of methanol were combined with 300 μl of water to homogenize 10 mg of frozen material for 15 min at 70 °C. The polar fraction was dried under vacuum, and the residue was derivatized for 120 min at 37 °C (in 40 μl of 20 mg ml^-1^ methoxyamine hydrochloride (Sigma-Aldrich, cat. no. 593-56-6) in pyridine followed by a 30 min treatment at 37 °C with 70 μl of *N*-methyl-*N*-(trimethylsilyl)trifluoroacetamide (MSTFA reagent; Macherey-Nagel, cat. no. 24589-78-4). An autosampler Gerstel Multi-Purpose system (Gerstel GmbH & Co. KG, Mülheim an der Ruhr, Germany) was used to inject 1 μl of the samples in splitless mode to a chromatograph coupled to a time-of-flight mass spectrometer system (Leco Pegasus HT TOF-MS; Leco Corp., St Joseph, MI, USA). Helium served as the carrier gas, maintained at a constant flow rate of 2 ml s⁻¹, and gas chromatography (GC) was carried out using a 30 m DB-35 capillary column (Agilent). The injector was set to 230 °C, while both the transfer line and ion source were maintained at 250 °C. The oven temperature started at 85 °C and was ramped up to 360 °C at a rate of 15 °C per minute. Mass spectra acquisition began after a 180-second solvent delay, with a scan speed of 20 scans per second over an m/z range of 70 to 600. Data analysis of chromatograms and mass spectra was performed using Chroma TOF 4.5 (LECO) and TagFinder 4.2 software. Based on a computation of the retention index with a variance of no more than 5%, metabolites were annotated and compared to the reference data from the Golm Metabolome Database (http://gmd.mpimp-golm.mpg.de; (40)). The results were statistically analyzed using Student’s t-test, using a threshold of P<0.05 between the samples of the control and treated plants (Supplemental Dataset).

### Sequence analysis of GLXI;2 from Col-0 and IP-Pal-0

The *GLXI;2* (AT1G11840) cDNAs of the *A. thaliana* accessions Col-0 and IP-Pal-0 were synthesized from RNA extracted from rosettes of plants growing under control conditions. Sequencing, performed by Macrogen Inc. (South Korea), covered the full coding region from the start codon (ATG) to the stop codon. The coding sequences (CDS) are available in GenBank, submission ID 2943379.

Protein sequences were derived from the CDS using the EMBOSS Transeq tool (https://www.ebi.ac.uk/Tools/st/emboss_transeq/). Sequence alignment was conducted with Clustal Omega v1.2.4 (https://www.ebi.ac.uk/Tools/msa/clustalo/), and multiple sequence alignment of GLXI homologs, as shown in Figure 6A, was performed using MEGA7.

### Promoter analysis

To investigate natural variation in the *GLXI;2* promoter between the Col-0 and IP-Pal-0 ecotypes, we obtained the upstream regulatory sequences of the *AT1G11840* gene using the “Pseudogenomes Download” tool, available at https://tools.1001genomes.org/. Specifically, we extracted the 1.5 kb region upstream of the start codon (ATG), spanning Chr1:3993657– 3996044. Cis-acting regulatory elements (CREs) within the promoter sequences were identified using the PlantCARE database (41). Visualization of the identified CREs was performed with TBtools (42), and comparative analysis of motif presence, absence, and positional differences between the two ecotypes was conducted using custom R scripts.

### Statistical analysis

P-values for Figures 1, 3, 4, 8, 9, and 10 were calculated using one-way ANOVA followed by Dunnett’s post hoc test to assess significance relative to the Col-0 control. For Figures 2 and 5, statistical differences were evaluated using unpaired t-tests. In Figure 10, principal component analysis (PCA) was performed using the prcomp function from the base R stats package, and results were visualized with the factoextra package.

**Figure 2.**
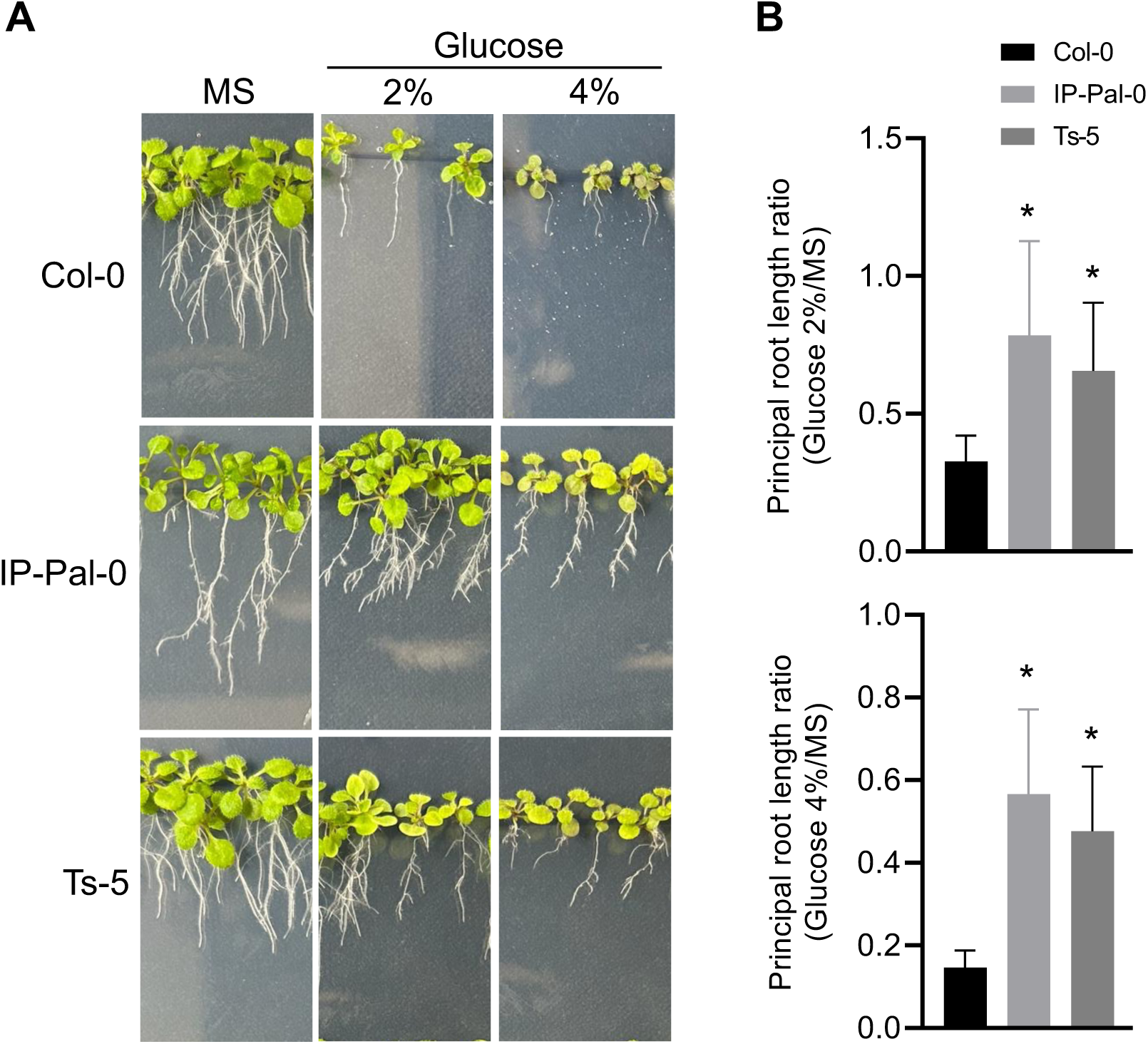
Development of seedlings of Col-0 and IP-PAL-0 in the presence of 2% and 4% glucose in the growth media. (A) Images of seedlings 14 d after germination in different growth media. Scale bar = 1 cm. (B) Main root length after 14 days of germination on media containing 2% or 4% glucose compared to the length in control media (MS). The graphs show the means and standard errors of the measurements of 3 independent experiments (*n*=3) of 20 roots each. An asterisk (*) indicates a significant difference (P <0.001) between Col-0 and IP-PAL-0 determined by unpaired t-test.

## Results

### Growth of *A. thaliana* ecotypes in the presence of reactive carbonyl species

To identify *A. thaliana* ecotypes with an enhanced capacity for RCS detoxification, we conducted a toxicity assay on 20 ecotypes from five distinct geographical regions (Fig. 1A). We examined plant development in the presence of 0.2 mM KDG or 0.5 mM MGO, concentrations previously established in studies on the Col-0 ecotype (8, 15). It is important to note that under our experimental conditions, 0.2 mM KDG does not significantly impair Col-0 development, while 0.5 mM MGO substantially affects its growth (8), providing a benchmark for evaluating ecotypic responses.

After 14 days of exposure to MGO, primary root length was significantly reduced in all ecotypes compared to the control condition (MS medium) (Fig. 1B and D). However, the extent of this reduction varied among ecotypes. IP-Pal-0 exhibited the highest tolerance, with only a 65% reduction in primary root length, followed by Ts-5. In contrast, Col-0 was the most sensitive, showing an 85% reduction. Notably, under MGO treatment, IP-Pal-0 also developed larger and greener rosettes compared to Col-0 (Fig. 1D). Ecotypic responses to 0.2 mM KDG after 14 days also varied. While most ecotypes displayed primary root growth that was similar to or lower than that of Col-0, both IP-Pal-0 and Ts-5 again exhibited significantly enhanced growth under KDG treatment (Fig. 1C and D). Notably, IP-Pal-0 showed a 50% higher primary root length ratio compared to Col-0, further emphasizing its superior resilience relative to the reference ecotype.

Our previous study demonstrated that glucose supplementation in growth media leads to KDG accumulation in *A. thaliana* tissues (8). KDG is a metabolic byproduct generated through glucose oxidation or the degradation of glycosylated proteins (7, 43, 44). Given its potential impact on plant physiology, we investigated the effect of glucose on the growth of Col-0, IP-Pal-0, and Ts-5. We found that IP-Pal-0 and Ts-5 developed significantly longer primary roots than Col-0 when grown in media supplemented with 2% and 4% glucose (Fig. 2). IP-Pal-0 also showed larger and greener rosettes than Col-0 and Ts-5. Notably, the magnitude of these growth enhancements correlated positively with glucose concentration, suggesting a dose-dependent effect. These results suggest that the growth inhibition observed in vivo in glucose-containing media may be due, at least in part, to the accumulation of KDG and possibly other RCS such as MGO derived from glucose metabolism.

### Transcriptional analysis of GLXI genes in *A. thaliana* ecotypes

The toxicity assay revealed that IP-Pal-0 and Ts-5 exhibit increased tolerance to KDG and, to a lesser extent, MGO. Since GLXI;2 specifically metabolizes KDG (8) and all GLXI isoforms participate in MGO detoxification (15), we further investigated the transcriptional regulation of *GLXI* genes in selected *A. thaliana* ecotypes displaying higher and lower RCS tolerance than Col-0. To this end, we performed qRT-PCR analysis on roots and shoots of seedlings grown with and without KDG in the growth medium.

Under control conditions (MS medium), *GLXI* expression levels were generally similar across all analyzed ecotypes, with a few exceptions: *GLXI;1* expression was significantly lower in Taz-0 compared to Col-0 in both roots and shoots; IP-Pal-0 and Ts-5 exhibited a twofold increase in *GLXI;1* expression in shoots compared to Col-0; and *GLXI;2* expression was approximately twofold higher in IP-Pal-0 and Ts-5 than in Col-0 in both roots and shoots (Fig. 3).

**Figure 3.**
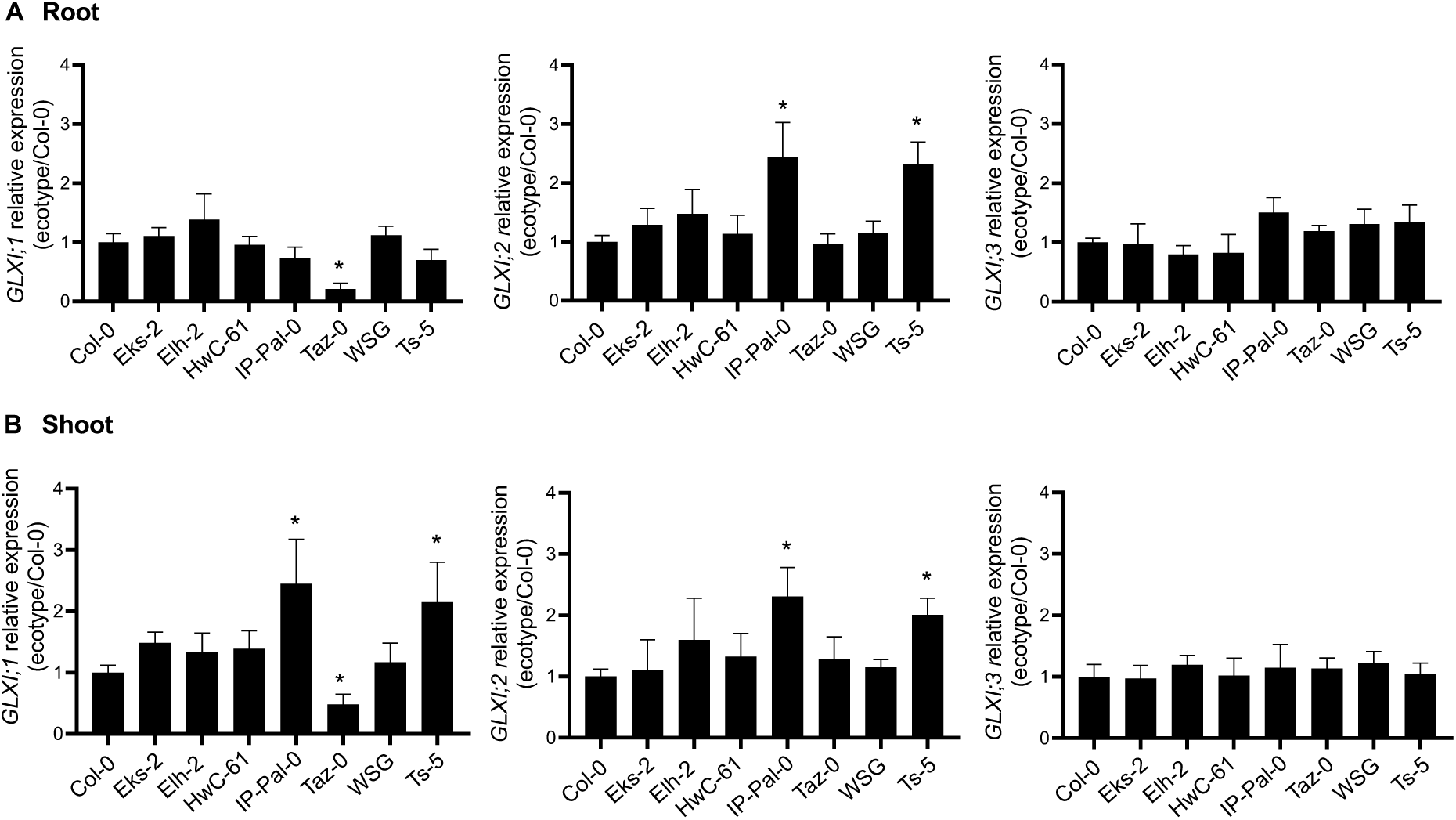
Transcriptional analysis of the GLXI isoforms in selected *A. thaliana* ecotypes grown on MS. Expression levels of GLXI isoforms in root (A) and shoot (B) of 14-week-old seedlings of selected ecotypes grown on MS relative to those measured in Col-0. One-way ANOVA was used to test for significant differences. An asterisk (*) indicates a significant difference (P<0.001) between an ecotype and Col-0 by Dunnett’s multiple comparison test.

Upon exposure to KDG, the expression of *GLXI;2* and *GLXI;3* was significantly higher in both roots and shoots of IP-Pal-0 and Ts-5, compared to Col-0 (Fig. 4). In IP-Pal-0 roots, the relative KDG/MS expression of *GLXI;2* increased by 3.7-fold, while *GLXI;3* was upregulated by 2.3-fold compared to Col-0 (Fig. 4A). Similarly, in IP-Pal-0 shoots, *GLXI;2* exhibited a striking 9.4-fold increase, and *GLXI;3* showed a 3.4-fold upregulation relative to Col-0 (Fig. 4B). Additionally, *GLXI;1* expression was upregulated in the shoots of IP-Pal-0 and Ts-5, with a 4-fold increase in IP-Pal-0 compared to Col-0 under KDG treatment. Other ecotypes displayed minor changes in *GLXI* isoform expression, but these variations were less pronounced than those observed in IP-Pal-0 and Ts-5.

**Figure 4.**
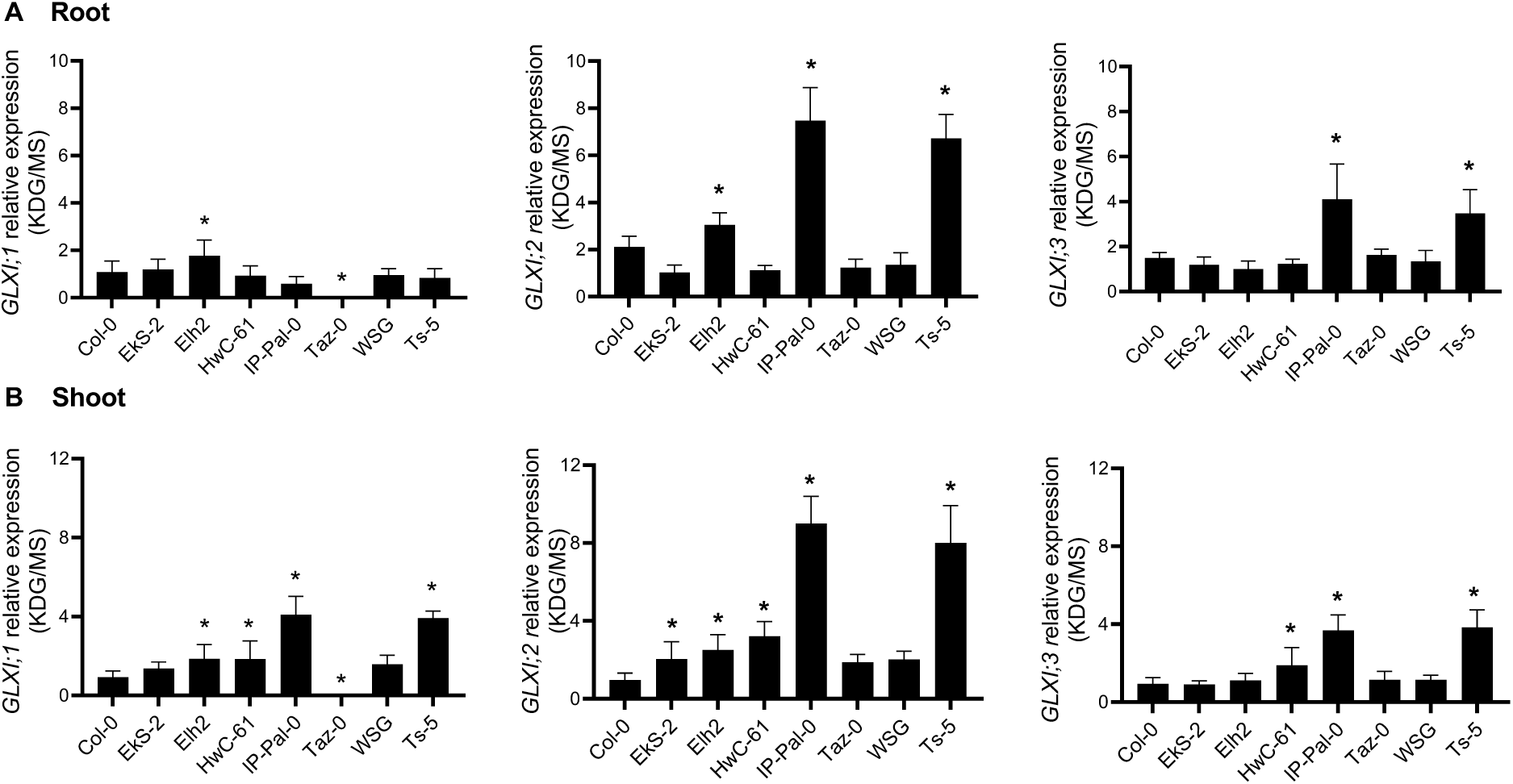
Transcriptional analysis of the GLXI isoforms in selected *A. thaliana* ecotypes grown on media with and without KDG. Expression levels of GLXI isoforms in root (A) and shoot (B) of 14-week-old seedlings of selected ecotypes grown in 0.2 mM KDG relative to those measured in seedlings grown in MS. One-way ANOVA was used to test for significant differences. An asterisk (*) indicates a significant difference (P<0.001) between an ecotype and Col-0 by Dunnett’s multiple comparison test.

Overall, qRT-PCR analysis demonstrated a substantial increase in *GLXI* transcription, particularly *GLXI;2*, in IP-Pal-0 and Ts-5 in response to high KDG concentrations.

### Metabolization of KDG by protein extracts of *A. thaliana* ecotypes

We further examined total GLXI activity in whole seedlings of selected *A. thaliana* ecotypes grown in the presence of 0.2 mM KDG. To assess KDG metabolization capacity, we measured free GSH concentrations after incubating protein extracts with varying KDG concentrations and a fixed GSH concentration, following our previously established protocol (8, 21). Our analysis revealed that IP-Pal-0 exhibited a statistically significant increase in KDG metabolization compared to Col-0, followed by Ts-5 (Fig. 5). These results further support the observed ecotypic differences in RCS detoxification and highlight IP-Pal-0 as the ecotypes with enhanced GLXI activity.

**Figure 5.**
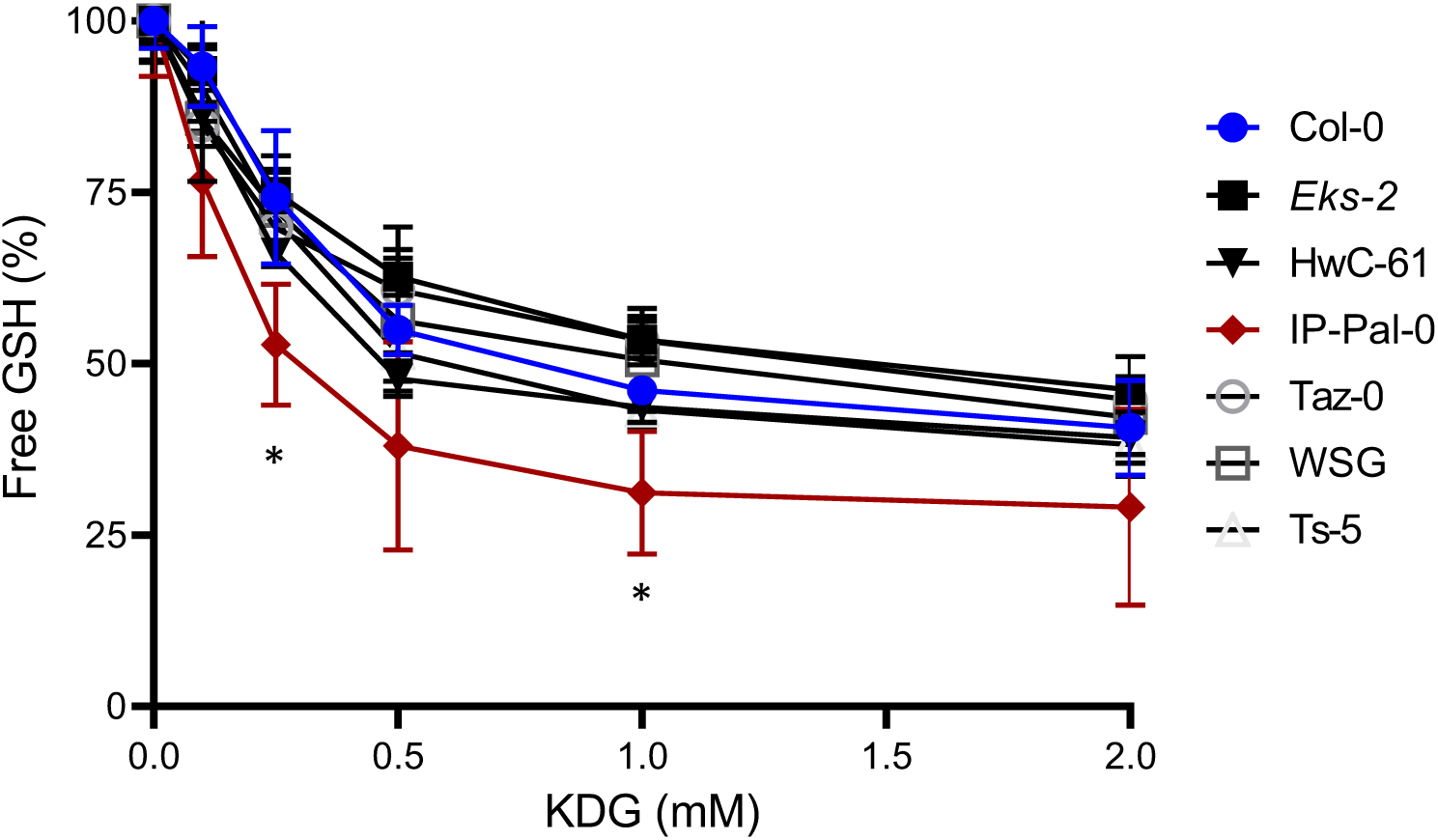
GLXI activity using KDG in protein extracts from seedlings of selected ecotypes. The concentration of free GSH was determined after incubation of protein extracts with different concentrations of KDG. The initial amount of free GSH when no KDG was added to the reaction media was assumed to be 100%. Values are mean ± SD (*n*=3). An asterisk (*) indicates a significant difference between ecotype and Col-0 values at each KDG concentration (P<0.05), determined by unpaired t-test.

### Comparison of GLXI;2 from different ecotypes

The higher GLXI activity observed in protein extracts from IP-Pal-0 compared to Col-0 is likely attributable to elevated GLXI;2 protein levels, consistent with the increased GLXI;2 transcript abundance detected in this ecotype (Fig. 3). However, we also considered the possibility that single nucleotide polymorphisms (SNPs) in the *GLXI;2* coding sequence of IP-Pal-0 could result in a more active enzyme variant. To investigate this, we analyzed the *GLXI;2* genomic sequence of IP-Pal-0 using data from the 1001 Genomes Project. Initial analysis revealed missing nucleotides in the IP-Pal-0 coding region. To clarify this discrepancy, we amplified and sequenced *GLXI;2* cDNA from leaf tissue of both Col-0 and IP-Pal-0 used in our study, identifying two notable SNPs. The first alters amino acid residue 229, with IP-Pal-0 encoding asparagine (Asn) instead of lysine (Lys) as found in Col-0 (Fig. 6A). The second affects residue 278, where IP-Pal-0 encodes isoleucine (Ile) in place of the phenylalanine (Phe) present in Col-0 (Fig. 6A).

**Figure 6.**
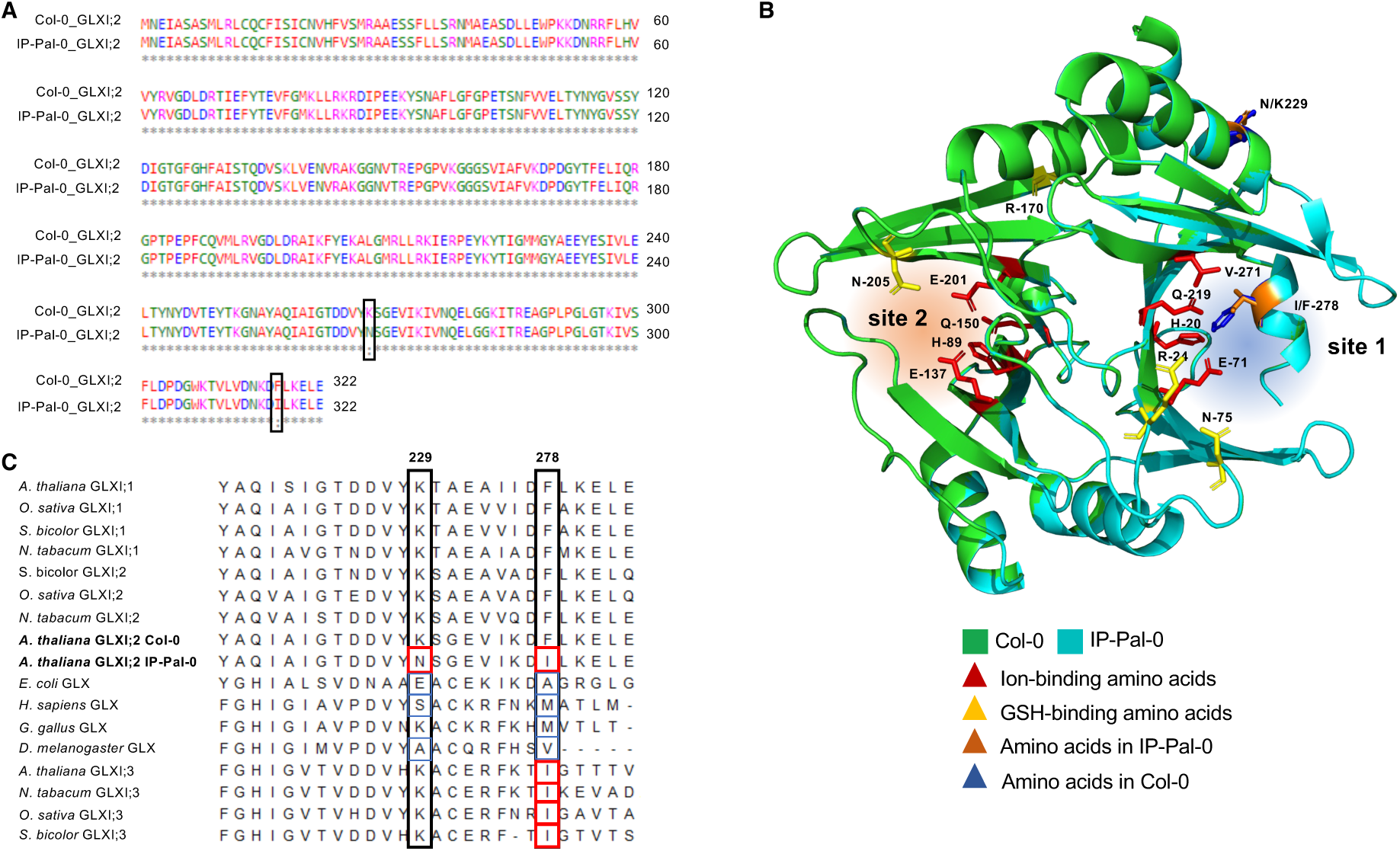
Analysis of GLXI;2 protein sequence and structure. (A) Alignment of the primary amino acid sequences of GLXI;2 from IP-Pal-0 and Col-0 highlighting the two amino acid differences between the ecotypes. (B) Model structures of Col-0 (green) and IP-Pal-0 (light blue) showing the localization of the amino acids at positions 229 and 278 and those involved in ion and GSH at the binding functional active site (site 2) and the equivalents in the cryptic site (site 1). (C) Alignment of selected Ni^2+^-type and Zn^2+^-type GLXI sequences from different organisms showing differential conservation of the amino acids at position 229 and 278 (corresponding to the *A. thaliana* Col-0 sequence). The analysis was conducted with the MEGA7 software. All sequences analysed are listed in Supplementary Table 1.

GLXI;2 is a monomeric enzyme with two distinct structural domains, forming two non-equivalent concavities. One of these, the cryptic site (site 1, Fig. 6B), cannot bind metal ions, whereas the other, site 2, serves as the functional metal-dependent active site (21). To assess the potential impact of the identified SNPs, we performed comparative structural modeling of GLXI;2 from Col-0 and IP-Pal-0, focusing on the localization of residues 229 and 278. Our analysis revealed that both residues are situated within structural domain B, contributing to the concavity of the cryptic site 1. Importantly, these residues are distant from the metal-ion binding residues in the active site (His89, Glu137, Gln150, and Glu201) (8, 21) (Fig. 6B).

To experimentally determine whether the Asn229 and/or Ile278 substitutions in IP-Pal-0 influence enzymatic function, we performed site-directed mutagenesis on Col-0 GLXI;2, generating K229N, F278I, and K229N-F278I mutants. As GLXI;2 activity cannot be spectrophotometrically measured using KDG as a substrate, we assessed the catalytic efficiency of the wild-type and mutant GLXI;2 variants using MGO and GO, along with their respective optimal metal cofactors (8, 15) (Table 1). Kinetic analysis, evaluating turnover rate (*k*_cat_), substrate affinity (*K*_m_), and catalytic efficiency (*k*_cat_/*K*_m_), yielded values consistent with previously published data (Schmitz et al., 2017). Importantly, we observed no significant differences between Col-0 GLXI;2 WT and the mutant variants, indicating that substitutions at residues 229 and 278 do not impact the catalytic efficiency of GLXI;2 under our assay conditions (Table 1). Additionally, native PAGE analysis showed no alterations in protein mobility among the different GLXI;2 variants (Suppl. Fig. S1), suggesting that these mutations do not affect the oligomeric organization of the enzyme.

**Table 1.** Kinetic parameters of Col-0 GLXI;2 WT and K229N, F278I, and K229N-F278I mutants using MGO and Ni^2+^, or GO and Mn^2+^. The kinetic constants were calculated by nonlinear regression analysis. Values are means ± SE (*n*=3) of independent enzyme preparations. Testing for significant differences was performed by t-test (P < 0.05).

| Col-0 GLXI;2 | MGO |  |  | GO |  |  |
| --- | --- | --- | --- | --- | --- | --- |
| | $K_m$<br>($\mu\text{M}$) | $k_{\text{cat}}$<br>($\text{s}^{-1}$) | $k_{\text{cat}}/K_m$<br>($\text{s}^{-1}\text{mM}^{-1}$) | $K_m$<br>($\mu\text{M}$) | $k_{\text{cat}}$<br>($\text{s}^{-1}$) | $k_{\text{cat}}/K_m$<br>($\text{s}^{-1}\text{mM}^{-1}$) |
| WT | 297 ± 14 | 7.5 ± 0.9 | 25.3 ± 1.7 | 411 ± 18 | 2.8 ± 0.3 | 6.8 ± 0.6 |
| K229N | 295 ± 23 | 9.1 ± 1.7 | 26.4 ± 2.2 | 415 ± 17 | 2.4 ± 0.3 | 6.5 ± 0.5 |
| F278I | 311 ± 18 | 8.4 ± 1.2 | 27.0 ± 2.4 | 403 ± 12 | 2.5 ± 0.4 | 6.2 ± 0.3 |
| K229N-F278I | 302 ± 15 | 8.1 ± 1.1 | 25.8 ± 2.1 | 407 ± 15 | 2.6 ± 0.3 | 6.7 ± 0.5 |

To determine whether the Asn229 and Ile278 residues in IP-Pal-0 GLXI;2 are conserved in other GLXI homologs, we conducted a comparative sequence analysis of GLXI proteins from diverse evolutionary lineages (Supplementary Table 1) (8). Notably, neither SNP was found in homologous positions within any other Viridiplantae-specific GLXI;1 or GLXI;2 isoforms (Fig. 6C; Supplementary Table 1). However, we identified a recurring pattern in Viridiplantae Zn²⁺-type GLXI enzymes, such as *A. thaliana* GLXI;3, where all sequenced enzymes contain an Ile at the position corresponding to residue 278 in IP-Pal-0 GLXI;2 (Fig. 6C; Supplementary Table 1). In contrast, GLXI enzymes from other organisms predominantly contain different amino acids at positions 229 and 278 (Fig. 6C; Supplementary Table 1).

### Analysis of *GLXI;2* promoter regions of different ecotypes

Since the enzymatic properties of GLXI;2 of Col-0 and IP-Pal-0 (Col-0 K229N-F278I) are very similar (Table 1), we propose that the high expression of *GLXI;2* observed in IP-Pal-0 in the presence of KDG is most likely responsible for the enhanced growth of this ecotype. To investigate this, we analyzed and compared the promoter regions of Col-0 and IP-Pal-0 for differences in cis-regulatory elements (CREs) and for conserved CREs located in distinct positions. Using the online database *Plant CARE*, we identified approximately 23 motifs present in both ecotypes (Fig. 7A) (Supplementary Table 2). Some motifs were found to share similar positions in both Col-0 and IP-Pal-0, while others were located in different positions within the region from −1250 to −800 (Supplementary Table 3). When we compared the sequences of the identified CREs between both ecotypes, we detected ecotype-specific motifs such as ERE (ATTTCATA) and MYC (CAATTG) which are only found in Col-0, and the ABRE (ACGTG; CACGTG; GCAACGTGTC) which is only found in IP-Pal-0 (Fig. 7B). This analysis suggests that specific motifs and *cis*-regulatory elements may be involved in the differences in *GLXI;2* gene expression related to KDG detoxification in Col-0 and IP-Pal-0.

**Figure 7.**
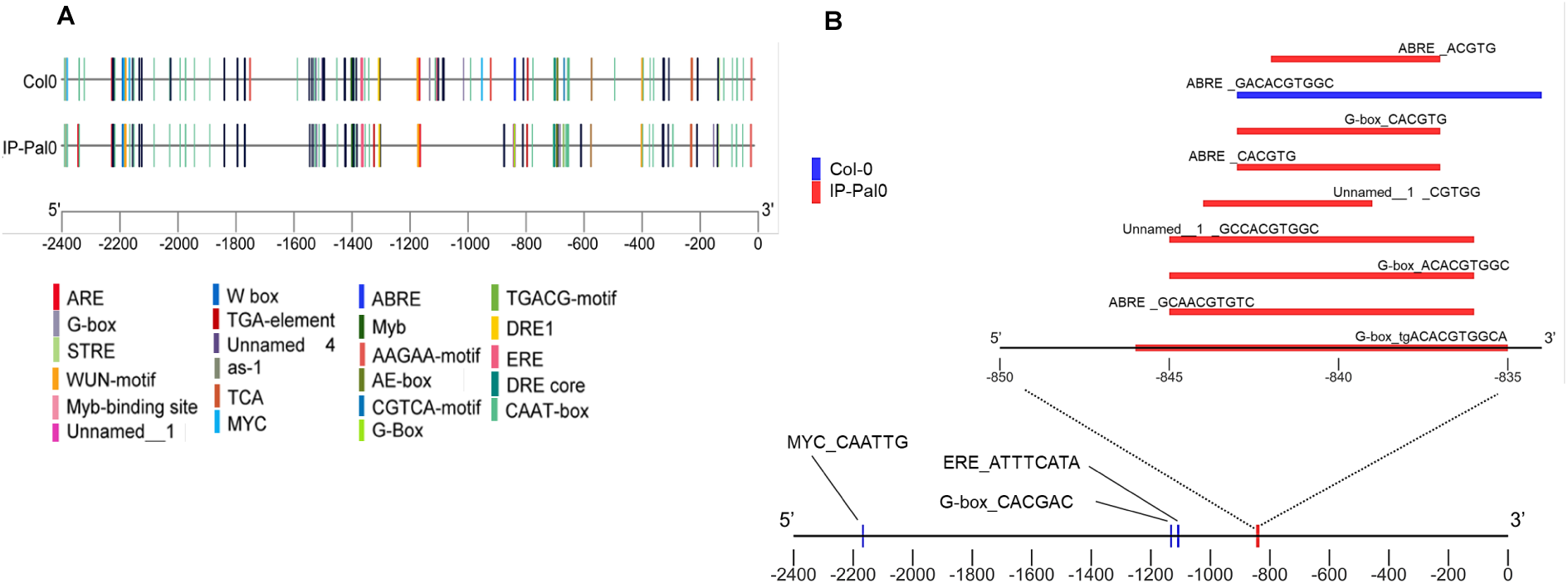
*Cis*-regulatory elements in Col-0 and IP-PAL-0. (A) Promoter analysis for the presence of cis-regulatory elements 1.5 kb upstream of the 5’ UTR (2.4 kb upstream of the transcription initiation site) of *GLX1;2* in the *A. thaliana* ecotypes Col-0 and IP-Pal-0. Cis-regulatory elements were mapped using TBtools (42). (B) Unique cis-regulatory elements of *GLX1;2* identified in the ecotypes Col-0 and IP-Pal-0.

### Transcriptional analysis of GLXII genes in *Arabidopsis thaliana* ecotypes

Building upon our previous findings that KDG metabolism in *A. thaliana* involves the sequential action of GLXI;2 and either GLXII;4 or GLXII;5 (8), we investigated whether the expression of these second-step genes is also modulated in response to KDG exposure. To this end, we performed qRT-PCR analysis of all *GLXII* genes in the same ecotypes previously examined for *GLXI* expression.

Under control conditions (MS medium), *GLXII* expression patterns were largely consistent across ecotypes, with a few notable differences. *GLXII;2* expression was approximately 0.5-fold lower in the roots of EkS-2 and Taz-0 compared to Col-0, whereas *GLXII;4* and *GLXII;5* expression was upregulated in the shoots of IP-Pal-0, showing a 0.6-fold increase relative to Col-0 (Fig. 8).

**Figure 8.**
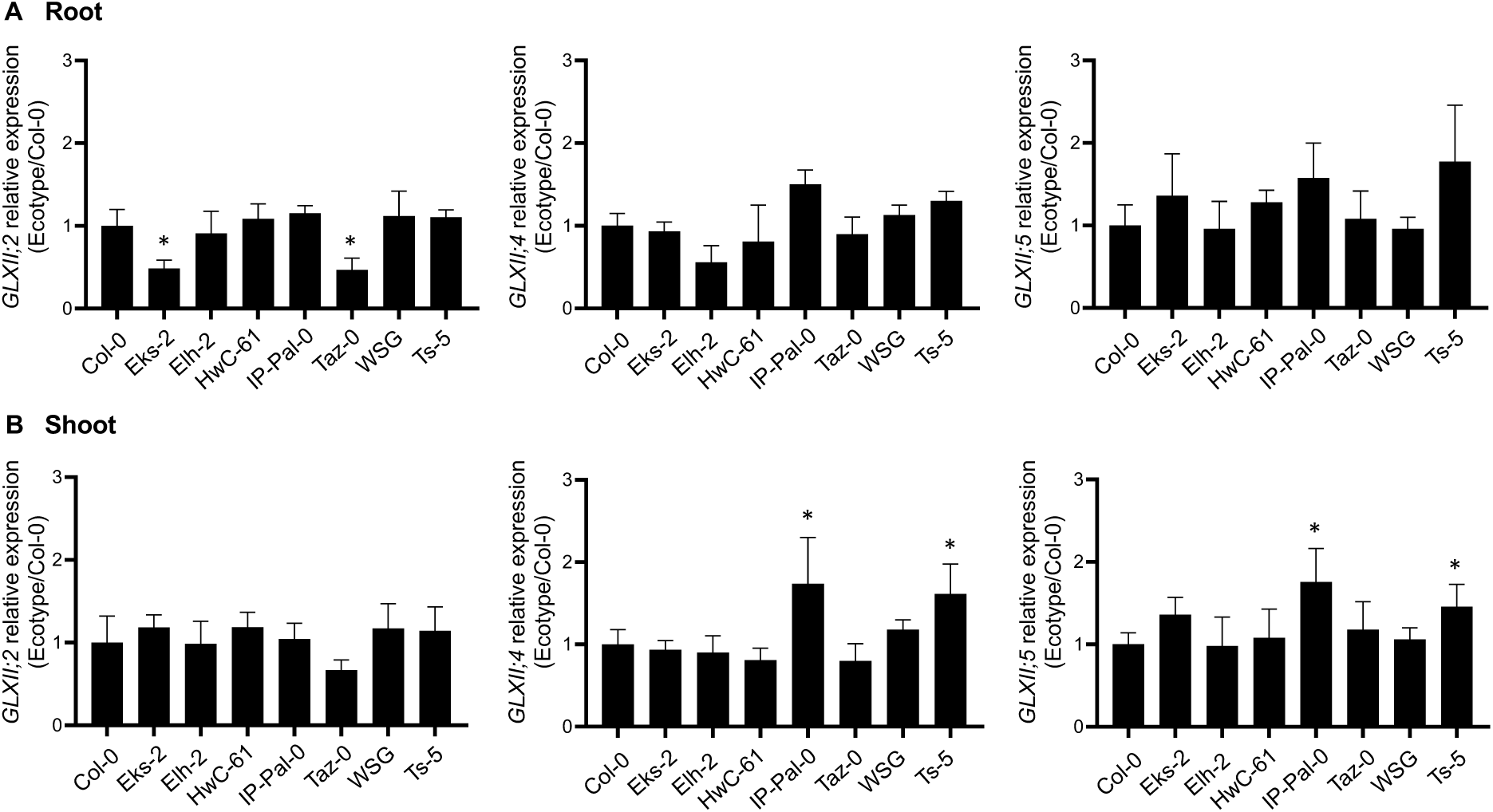
Transcriptional analysis of the GLXII isoforms in selected *A. thaliana* ecotypes grown on MS. Expression levels of GLXII isoforms in root (A) and shoot (B) of 14-week-old seedlings of selected ecotypes grown on MS relative to those measured in Col-0. One-way ANOVA was used to test for significant differences (P<0.0001). An asterisk (*) indicates a significant difference (P<0.001) between an ecotype and Col-0 by Dunnett’s multiple comparison test.

Upon KDG exposure, distinct isoform-specific expression patterns emerged. In IP-Pal-0 roots, *GLXII;2* expression remained unchanged compared to Col-0, while *GLXII;4* and *GLXII;5* expression showed a dramatic increase, with a 13-fold and 12-fold upregulation, respectively (Fig. 9A). In shoots, *GLXII;2* and *GLXII;5* were expressed at levels 0.5-fold higher than in Col-0, while *GLXII;4* expression was 3.3-fold higher (Fig. 9B). Notably, Ts-5 exhibited a transcriptional response highly similar to that of IP-Pal-0. Other ecotypes also displayed specific changes in GLXII isoform expression, often consistently across both roots and shoots (Fig. 9B).

**Figure 9.**
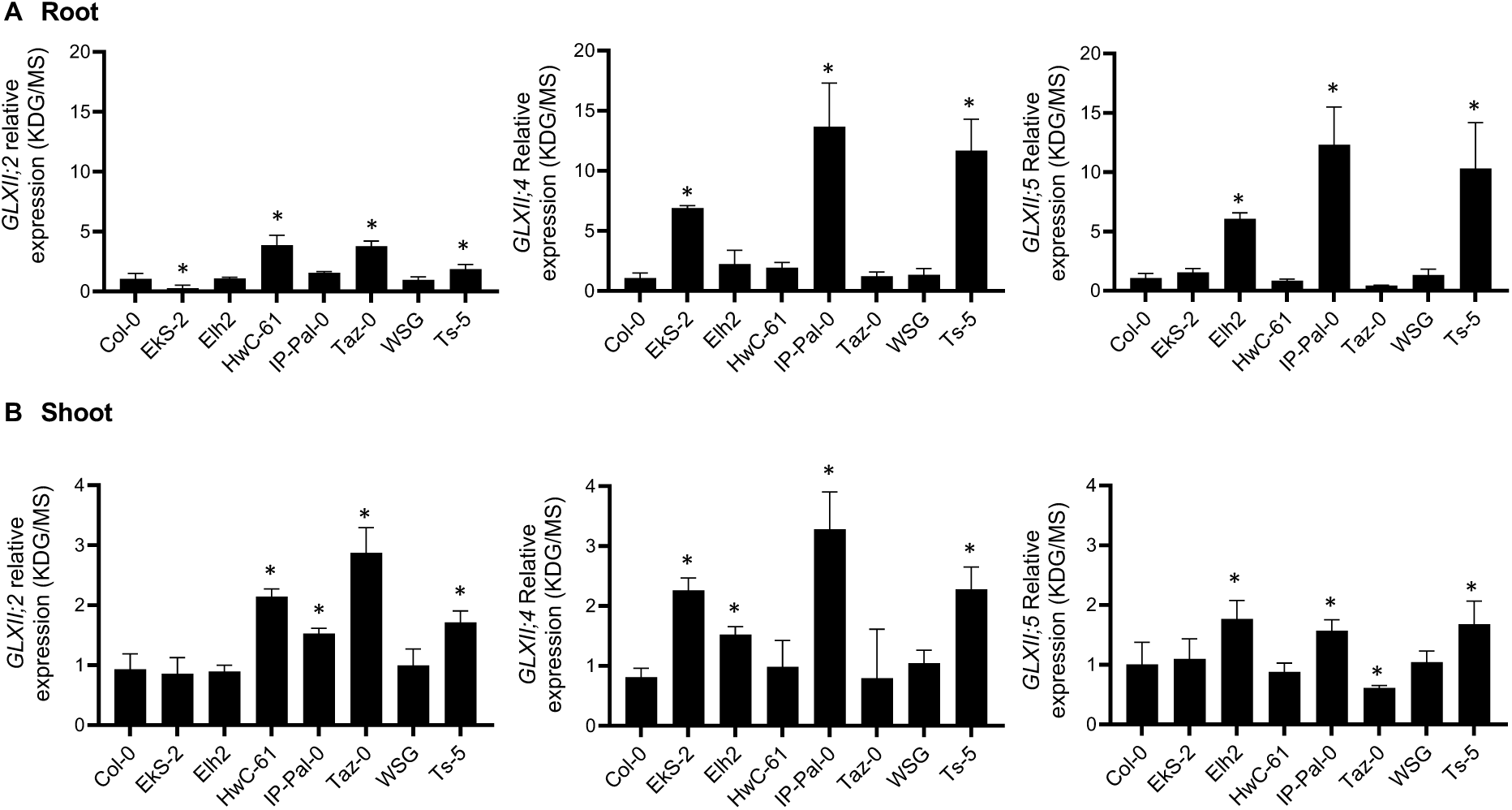
Transcriptional analysis of the GLXII isoforms in selected *A. thaliana* ecotypes grown on media with and without KDG. Expression levels of GLXI isoforms in root (A) and shoot (B) of 14-week-old seedlings of selected ecotypes grown in 0.2 mM KDG relative to those measured in seedlings grown in MS. One-way ANOVA was used to test for significant differences (P<0.0001). An asterisk (*) indicates a significant difference (P<0.001) between an ecotype and Col-0 by Dunnett’s multiple comparison test.

These results, in line with our previous findings (8), underscore the role of *GLXII;4* and *GLXII;5* in the second step of KDG detoxification, particularly in IP-Pal-0 and Ts-5.

### Metabolite profile analysis in seedlings of Col-0 and IP-Pal-0 grown in MS and KDG

We hypothesized that if IP-Pal-0 is more effective than Col-0 at detoxifying KDG, the two ecotypes would exhibit distinct metabolic profiles. To test this hypothesis, we performed a metabolite profiling analysis via GC-MS on seedling extracts of IP-Pal-0 and Col-0 grown in 0.2 mM KDG and MS control medium (Supplementary Data 1). To identify patterns and groupings within the dataset, we performed principal component analysis (PCA) (Fig. 10A and Suppl. Fig. S2). The first principal component (PC1) explained 44.6% of the variance, effectively separating Col-0 from IP-Pal-0, while the second principal component (PC2) explained 22.3% of the variance, distinguishing the different growth conditions.

**Figure 10.**
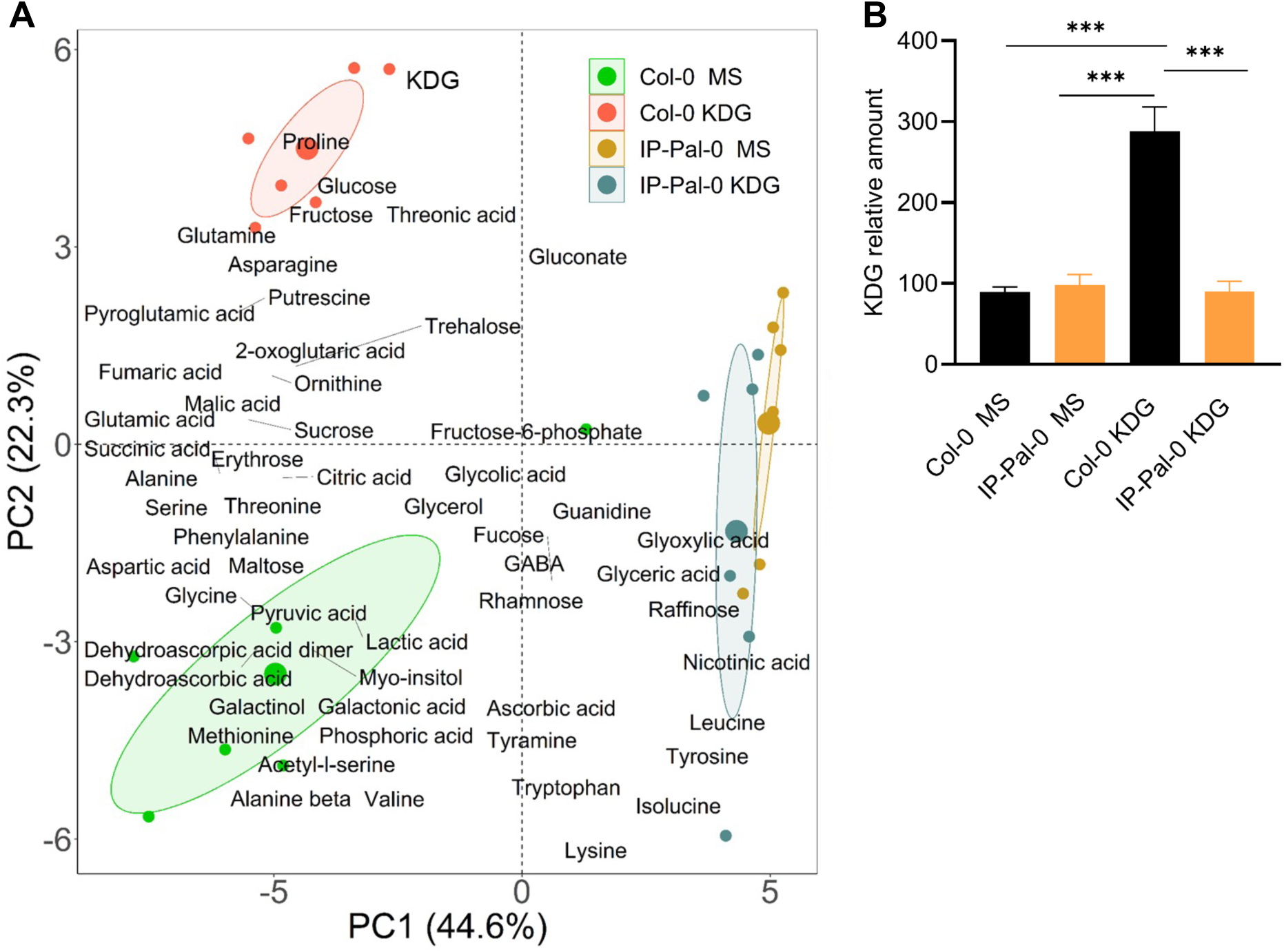
Analysis of the metabolite relative contents. (A) 2D Principal Component Analysis (PCA) biplot of variables in black labels (metabolites measured in seedlings grown in media without (MS) or with 0.2 mM KDG and individuals in colored points (*A. thaliana* ecotypes Col-0 and IP-Pal-0). Ellipses group the ecotypes per growth medium. (B) Relative amount of KDG in seedlings of Col-0 and IP-Pal-0 grown in media without (MS) and with 0.2 mM KDG. Barplots with one asterisk or more are significantly different (Tukey’s HSD, P < 0.1; *P < 0.05; **P < 0.01; ***P < 0.001).For statistical analysis multiple comparison tests using Tukey’s honest significant difference test (Tukey HSD) were used to compare the barplots.

Notably, the PCA revealed a distinct separation between Col-0 grown in MS and KDG-supplemented conditions, suggesting significant metabolic changes in response to KDG. In contrast, IP-Pal-0 grown in MS and KDG exhibited similar relative metabolite levels, indicating more stable metabolic profile regardless of KDG exposure. The PCA biplot, which includes loadings (Fig. 10A), highlights metabolites that strongly contribute to PC variation. KDG, along with sugars such as fructose and glucose, as well as proline, exhibited high loadings on PC2, particularly in the direction of Col-0 grown in KDG-containing media, suggesting that Col-0 is experiencing physiological stress under these conditions.

Next, we examined the relative KDG content in Col-0 and IP-Pal-0 obtained by GC-MS (Fig. 10B). We found that Col-0 grown in the presence of KDG accumulated about 2.5-fold higher amounts of KDG than the plants grown in MS. Interestingly, IP-Pal-0 grown in MS or in the presence of KDG showed no differences in relative KDG levels, suggesting an enhanced capacity for the detoxification of this RCS.

## Discussion

Understanding how natural variation shapes stress tolerance is crucial for both basic plant biology and future crop improvement. Our findings reveal that a specific *A. thaliana* ecotype, IP-Pal-0, has evolved an enhanced capacity to detoxify glucose-derived RCS, highlighting a previously underappreciated stress defense mechanism. Given that glucose-derived RCS such as KDG accumulate during high-sugar fluxes triggered by abiotic stress like drought or salinity, the enhanced detoxification capacity of IP-Pal-0 likely represents a naturally selected trait for survival in oxidative environments.

The *A. thaliana* ecotypes studied in this work originate from different geographical regions (Fig. 1A) and may therefore have been subject to different selective pressures. As a result, the ecotypes may have developed adaptive responses to environmental stresses, which we have exploited here to identify ecotypes with an enhanced capacity for glucose-derived RCS detoxification. After growing the ecotypes in the presence of MGO and KDG, which are specifically produced under oxidative stress, the IP-Pal-0 ecotype showed the highest tolerance to the toxic effects of both RCS, especially KDG, compared to most of the other ecotypes (Fig 1B-D). A second ecotype, Ts-5, also showed a similar behavior, but to a much lesser extent. It is interesting to note that both ecotypes originate from very restricted specific regions in Spain (Fig. 1A).

Here we show that in IP-Pal-0, *GLXI;2*, *GLXII;4* and *GLXII;5* are highly expressed in roots and shoots when plants are grown in the presence of RCS, particularly KDG (Fig. 4 and Fig. 8). In line with these elevated expression levels, protein extracts from IP-Pal-0 showed significantly higher total GLXI activity using KDG as a substrate compared to extracts from other ecotypes (Fig. 5). In the Ts-5 ecotype, we observed increased expression of *GLXI;2*, *GLXII;4*, and *GLXII;5*, though to a lesser extent than in IP-Pal-0. Interestingly, despite this upregulation of expression, whole-seedling protein extracts from Ts-5 displayed GLXI activity with KDG similar to that of Col-0. We hypothesize that using entire seedlings may obscure tissue-specific differences in enzymatic activity, making the assay insufficiently sensitive to detect subtle variations that could be present in individual tissues.

We further found that IP-Pal-0 does not accumulate KDG when grown on media containing this RCS, whereas Col-0 does (Fig. 10B). These results suggest that the expression of *GLXI;2* and *GLXII;4* and *GLXII;5* is enhanced by KDG, although it cannot be excluded that the high level of KDG in the growth media further increases the production of other RCS and their derivatives, and thus the plant also responds to them by increasing its scavenging capacity. It is worth mentioning that although GLXI;2 is specifically involved in the detoxification of KDG (8), it can also act on the hemithioacetal formed by MGO and GSH (15). In any case, the high expression levels of GLXI;2 and GLXII;4 and GLXII;5 in response to KDG in the growth media are consistent with previous work showing that these isoforms are specifically involved in the detoxification of KDG (8).

In addition to the high expression of GLXI;2 in IP-Pal-0, another possibility is that this isoform may be more active than the Col-0 isoform. Upon examining the sequence of IP-Pal-0 GLXI;2, we identified two different amino acids compared to Col-0. We produced single and double mutants of the Col-0 protein to match those changes in IP-Pal-0 but found no differences in the basic kinetic behavior between the two proteins (Table 1). Mapping the mutations in the modelled structure of GLXI;2 revealed that both residues are located far away from the active site in the part of domain B involved in the formation of the cryptic site (Fig. 6B). It has been proposed that this cryptic site has lost its catalytic capacity due to the accumulation of mutations that do not allow the binding of the metal required for catalysis (21, 23). Interestingly, we found that the specific combination of amino acids found in IP-Pal-0 GLXI;2 (Asn instead of Lys 229, and Ile instead of Phe 278) does not occur in GLXI proteins of any other organism, but that all sequenced Viridiplantae Zn^2+^-type GLXI contain the Ile at position 278 (Fig. 6C; Supplementary Table 1). The recurrent presence of Ile at position 278 in distantly related plant GLXI enzymes may reflect convergent evolution toward a structurally or functionally advantageous configuration, even though this residue lies outside the active site. In contrast to Ni^2+^-type GLXI, Viridiplantae Zn^2+^-type GLXI are monomeric proteins with two structural domains that form two similar active sites (8), so the presence of Ile at this position does not appear to be detrimental to the catalytic function. It is also possible that post-translational modifications in IP-Pal-0 contribute to altered GLXI;2 protein properties. This presents an interesting avenue for future investigation. Importantly, we found that specific motifs and *cis*-regulatory elements within the Col-0 and IP-Pal-0 GLXI;2 promoter regions may be involved in the differences in GLXI;2 gene expression associated with KDG detoxification. In the future, this knowledge could be used to identify important promoter regions for the enhancement of this trait.

## Conclusions

Our findings highlighted the significant potential of naturally occurring *A. thaliana* variants as valuable genetic resources for improving plant resilience under diverse environmental conditions. Specifically, glucose-derived RCS such as KDG and MGO, accumulate in response to drought, salinity, high temperature, and other oxidative stresses; identifying *GLXI;2*, *GLXII;4*, and *GLXII;5* as key detoxification components underscores the crucial role of the glyoxalase system in alleviating cellular damage. By exploiting high-expression alleles or stress-inducible promoters from tolerant ecotypes (e.g., IP-Pal-0), breeders and biotechnologists can potentially increase stress tolerance in crops without the need for transgenic methods. This approach aligns with current trends in agricultural regulation and public acceptance.

Furthermore, identifying naturally occurring regulatory elements in the *GLXI;2* promoter paves the way for cis-regulatory engineering to optimize detoxification pathways. Given that glyoxalase-mediated RCS scavenging plays a critical role in plant metabolism and defense, targeted modulation of its components could improve performance across diverse growing environments. More broadly, these findings emphasize the power of leveraging natural genetic diversity - particularly within well-characterized model species - to uncover adaptive strategies for overcoming environmental challenges, ultimately guiding the development of more robust and climate-resilient crops.

## Supplementary material

**Supplementary Figure 1.** Native PAGE of GLXI;2. Two μg of recombinant GLXI;2 of Col-0 WT, and the mutants K229N, F278I, K229N-F278I were analysed in a 10% (w/v) native PAGE and visualized with Coomassie Brilliant blue. MWM: SERVA Marker Mix for Blue/Clear Native PAGE (SERVA Electrophoresis GmbH; Heidelberg, Germany).

**Supplementary Figure 2.** Scree plot showing the variance explained by each principal component in the PCA. The plot aids in determining the number of components to retain for subsequent analysis.

**Supplementary Data 1. Relative amounts of metabolites determined by GC-MS.** Six biological replicates (BR1-BR6) were analyzed from each line.

**Supplementary Table 1. GLXI sequences used for the analysis of the amino acids at positions 229 and 278 corresponding to the *A. thaliana* GLXI;2 Col-0 sequence.** The sequences were extracted from the evolutionary analysis of GLXI presented in Supplementary Fig. 2 of Balparda et al. 2023.

**Supplementary Table 2. Cis-regulatory elements identified in Col-0 and IP-Pal-0**. The PlantCARE (41) tool was used. The positions are from 0 to −2400 bp in the 5’ direction (0 is in the 3’ end, which is the position of the last nucleotide before the ATG, and −2400 is in the 5’ end).

**Supplementary Table 3. Differences in the position of the identified cis-regulatory elements in Col-0 and IP-Pal-0.** The positions are from 0 to −2400 bp in the 5’ direction (0 is in the 3’ end which is the position of the last nucleotide before the ATG and −2400 is in the 5’ end).

## Declarations

### Ethics approval and consent to participate

Not applicable.

### Consent for publication

Not applicable.

### Availability of data and materials

All data supporting the findings of this study are provided in the main text and Supplementary Information. The GLXI;2 (AT1G11840) cDNA sequences from Col-0 and IP-Pal-0, covering the full coding region from start to stop codon, have been deposited in GenBank under submission ID 2943379. All other AT1G11840 sequences used for promoter analysis of the selected ecotypes were obtained from the 1001 Genomes Project via the “Pseudogenomes Download” tool, available at https://tools.1001genomes.org/.

### Authoŕs contributions

V.G.M. conceived and designed research. V.G.M., M. Bo and M.Ba. supervised data analysis and interpretation. M.Ba., A.K., Y.F., C.E.A., and S.A. performed research and acquired the raw data. A.R.F. contributed with analytical tools. V.G.M., M.Ba., and M.Bo. wrote the manuscript. All authors contributed to the generation of the figures and the revision of the manuscript. All authors accepted the final version of the manuscript.

