## Supplementary material for "Natural variation in *Arabidopsis thaliana* highlights a key role of Glyoxalase I;2 in detoxifying glucose-derived reactive carbonyl species": Suppl Table 1. docx.docx

**Supplementary Table 1. GLXI sequences used for the analysis of amino acid residues at positions 229 and 278, based on the *A. thaliana* GLXI;2 Col-0 reference sequence.** Sequences for Col-0 and IP-Pal-0 were obtained in this study by sequencing GLXI;2 cDNA from leaf tissue of both accessions. All other sequences were sourced from the evolutionary analysis of GLXI presented in Supplementary Figure 2 of Balparda et al. (2023).

| **Abbreviation** | **NCBI GLX corresponding sequence code** | **229** | **278** |
| --- | --- | --- | --- |
| *A. thaliana* GLXI;1 | AAM61701.1 *Arabidopsis thaliana* Eudicot | K | F |
| *E. salsugineum* GLXI;1 | XP 006391510.1 *Eutrema salsugineum* Eudicot | K | F |
| *O. sativa* GLXI;1 | BAD28547.1 *Oryza sativa* Japonica Monocot | K | F |
| *S. bicolor* GLXI;1 | XP 021315635.1 *Sorghum bicolor* Monocot | K | F |
| *L. japonicus* GLXI;1 | AFK49540.1 *Lotus japonicus* Eudicot | K | F |
| *E. guineensis* GLXI;1 | XP 010915890.1 *Elaeis guineensis* Monocot | **R** | F |
| *V. vinífera* GLXI;1 | XP 002283968.1 *Vitis vinifera* Eudicot | K | F |
| *N. tabacum* GLXI;1 | XP 016444062.1 *Nicotiana tabacum* Eudicot | K | F |
| *C. variabilis* GLXI;2 | XP 005849034.1 *Chlorella variabilis* Chlorophyta | K | F |
| *C. subellipsoidea* GLXI;2 | XP 005648190.1 *Coccomyxa subellipsoidea* Chlorophyta | K | F |
| *E. guineensis* GLXI;2 | XP 010931859.1 *Elaeis guineensis* Monocot | K | F |
| *S. bicolor* GLXI;2 | XP 021321462.1 *Sorghum bicolor* Monocot | K | F |
| *O. sativa* GLXI;2 | BAB71741.1 *Oryza sativa* Japonica Group Monocot | K | F |
| *L. japonicus* GLXI;2 | AFK48160.1 *Lotus japonicus* Eudicot | K | F |
| *N. tabacum* GLXI;2 | XP 016442481.1 *Nicotiana tabacum* Eudicot | K | F |
| *V. vinifera* GLXI;2 | XP 002273346.2 *Vitis vinífera* Eudicot | K | F |
| ***A. thaliana* GLXI;2 Col-0** | ***Arabidopsis thaliana* Eudicot** | K | F |
| ***A. thaliana* GLXI;2 IP-Pal-0** | ***Arabidopsis thaliana* IP-Pal-0 Eudicot** | **N** | **I** |
| *E. salsugineum* GLXI;2 | XP 006417306.1 *Eutrema salsugineum* Eudicot | K | F |
| *S. moellendorffii* GLXI;2 | XP 002960285.1 *Selaginella moellendorffii* Lycopodiophyta | K | F |
| *P. patens* GLXI | PNR26300.1 *Physcomitrella patens* Bryophyta | K | F |
| *N. mirabilis* GLXI | *GBST01006244.1 Nitella mirabilis* Charophyta | K | F |
| *C. fungivorans* GLXI | WP 061541456.1 *Collimonas fungivorans* Bacteria | K | **G** |
| *S. aquatica* GLXI | WP 133329806.1 *Sapientia aquatica* Bacteria | **E** | **D** |
| *E. coli* GLXI | BAE76494.1 *Escherichia coli* GLXI Bacteria | **E** | **A** |
| *S. boydii* GLXI | WP 072117963.1 *Shigella boydii* Bacteria | **E** | **A** |
| *C. amalonaticus* GLXI | WP 061070167.1 *Citrobacter amalonaticus* Bacteria | **E** | **A** |
| *K. sacchari* GLXI | WP 017456599.1 *Kosakonia sacchari* Bacteria | **Q** | **A** |
| *S. bongori* GLXI | WP 001237803.1 *Salmonella bongori* Bacteria | **E** | **A** |
| *R. chamberiensis* GLXI | WP 072045048.1 *Rouxiella chamberiensis* Bacteria | **Q** | **V** |
| *P. fontium* GLXI | WP 074822919.1 *Pragia fontium* Bacteria | **A** | **A** |
| *B. diplopodorum* GLXI | WP 159566424.1 *Budvicia diplopodorum* Bacteria | **Q** | **A** |
| *C. apiculatus* GLXI | WP 044249178.1 *Chondromyces apiculatus* Bacteria | **G** | **R** |
| *H. ochraceum* GLXI | WP 012831609.1 *Haliangium ochraceum* Bacteria | K | **R** |
| *P. fumosum* GLXI | WP 136929061.1 *Polyangium fumosum* Bacteria | K | **E** |
| Cyanothece sp. CCY0110 GLXI | WP 008274325.1 Cyanothece sp. CCY0110 Cyanobacteria | **G** | **Q** |
| *N. punctiforme* GLXI | WP 012409889.1 *Nostoc punctiforme* Cyanobacteria | **A** | **Q** |
| *C. merolae* GLXI | XP 005535468.1 *Cyanidioschyzon merolae* Rhodophyta | **A** | **S** |
| *G. sulphuraria* GLXI | XP 005708421.1 *Galdieria sulphuraria* Rhodophyta | **R** | **S** |
| *A. flavus* GLXI | KOC17366.1 *Aspergillus flavus* Fungi | **A** | **L** |
| *P. triticirepentis* GLXI | XP 001937039.1 *Pyrenophora triticirepentis* Fungi | **S** | **L** |
| *C. gattii* GLXI | KIR81401.1 *Cryptococcus gattii* Fungi | **A** | **R** |
| *S. cerevisiae* GLXI | EDN64389.1 *Saccharomyces cerevisiae* Fungi | **A** | **I** |
| *A. flavus* GLXI | KOC17366.1 *Aspergillus flavus* Fungi | **A** | **L** |
| *E. siliculosus* GLXI | CBJ26592.1 *Ectocarpus siliculosus* Phaeophyta | **G** | **I** |
| *T. gondii* GLXI | XP 018634997.1 *Toxoplasma gondii* Protozoa | **G** | F |
| *E. gracilis* GLXI | GDJR01027134.1 *Euglena gracilis* Protozoa | **A** | **Y** |
| *P. tricornutum* GLXI | XP 002180432.1 *Phaeodactylum tricornutum* Diatoms | K | F |
| *T. gondii* GLXI | XP 018634997.1 *Toxoplasma gondii* Protozoa | **G** | F |
| *E. siliculosus* GLXI | CBN74521.1 *Ectocarpus siliculosus* Phaeophyta | **G** | **M** |
| *A. aureolatum* GLXI | JAT91363.1 *Amblyomma aureolatum* Animalia | **A** | **V** |
| *M. musculus* GLXI | AAH24663.1 *Mus musculus* Animalia | **A** | **I** |
| *R. norvegicus* GLXI | NP 997477.1 *Rattus norvegicus* Animalia | **E** | **M** |
| *H. sapiens* GLXI | AAD38008.1 *Homo sapiens* Animalia | **S** | **M** |
| *D. rerio* GLXI | NP 998316.1 *Danio rerio* Animalia | **A** | **M** |
| *X. tropicalis* GLXI | NP 001025545.1 *Xenopus tropicalis* Animalia | **A** | **M** |
| *G. gallus* GLXI | XP 419481.1 *G. gallus* Animalia | K | **M** |
| *D. melanogaster* GLXI | NP 610270.1 *Drosophila melanogaster* Animalia | **A** | **V** |
| *S. kowalevskii* GLXI | XP 002737169.1 *Saccoglossus kowalevskii* Animalia | **A** | **M** |
| *A. queenslandica* GLXI | XP 003388487.1 *Amphimedon queenslandica* Animalia | K | K |
| *C. reinhardtii* GLXI | XP 001698653.1 *Chlamydomonas reinhardtii* Chlorophyta | **A** | **A** |
| *V. carteri* GLXI | XP 002956590.1 *Volvox carteri* f. nagariensis Chlorophyta | **A** | **S** |
| *N. mirabilis* GLXI;3 | GBST01009973.1 *Nitella mirabilis* Charophyta | K | **M** |
| *A. thaliana* GLXI;3 | NP 172291.1 *Arabidopsis thaliana* Eudicot | K | **I** |
| *E. salsugineum* GLXI;3 | XP 006417726.1 *Eutrema salsugineum* Eudicot | K | **I** |
| *N. tabacum* GLXI;3 | XP 016438641.1 *Nicotiana tabacum* Eudicot | K | **I** |
| *O. sativa* GLXI;3 | BAF17027.1 *Oryza sativa* Japonica Group Monocot | K | **I** |
| *S. bicolor* GLXI;3 | XP 021318420.1 *Sorghum bicolor* Monocot | K | **I** |
| *E. guineensis* GLXI;3 | XP 010943402.1 *Elaeis guineensis* Monocot | K | **I** |
| *L. japonicus* GLXI;3 | AFK46085.1 *Lotus japonicus* Eudicot | K | **I** |
| *V. vinífera* GLXI;3 | XP 003632261.1 *Vitis vinífera* Eudicot | K | **I** |
