## Supplementary figures and images for "Natural variation in *Arabidopsis thaliana* highlights a key role of Glyoxalase I;2 in detoxifying glucose-derived reactive carbonyl species"

### Suppl Fig S1.pdf

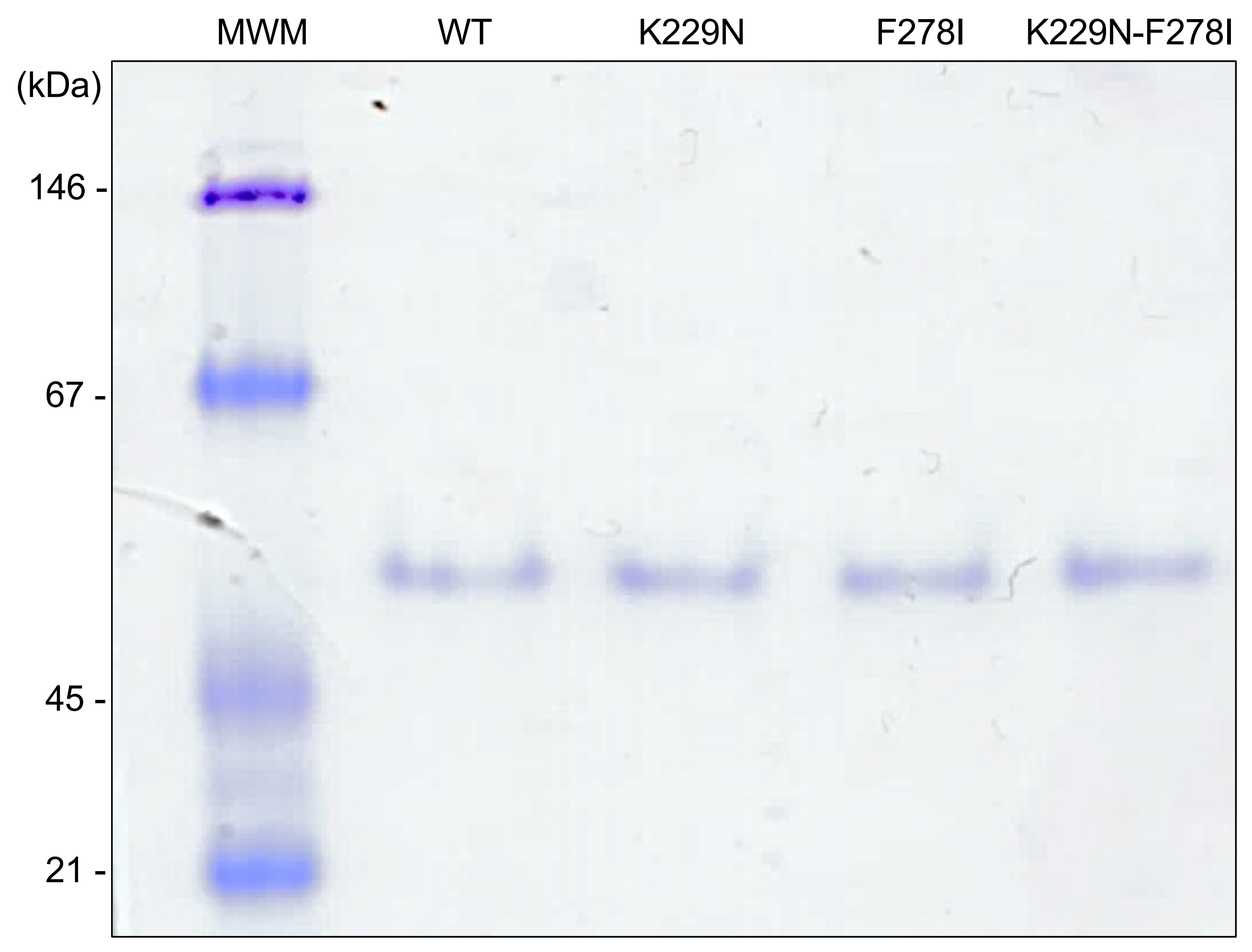

### Suppl Fig S2.pdf

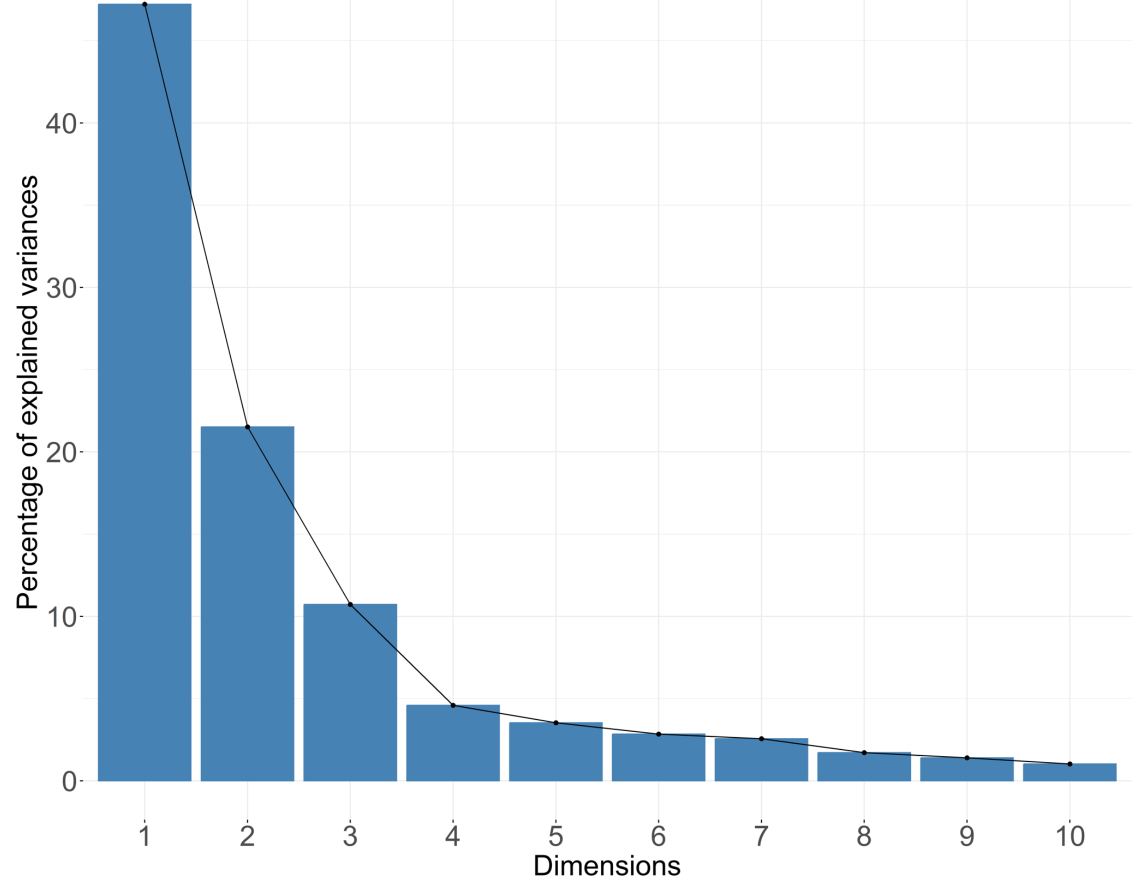
